# Acetaminophen production in the edible, filamentous cyanobacterium *Arthrospira platensis*

**DOI:** 10.1101/2022.06.30.498297

**Authors:** Jacob M. Hilzinger, Skyler Friedline, Divya Sivanandan, Ya-Fang Cheng, Shunsuke Yamazaki, Douglas S. Clark, Jeffrey M. Skerker, Adam P. Arkin

## Abstract

Spirulina is the common name for the edible, non-heterocystous, filamentous cyanobacterium *Arthrospira platensis* that is grown industrially as a food supplement, animal feedstock, and pigment source. Although there are many applications for engineering this organism^1–3^, until recently no genetic tools or reproducible transformation methods have been published. While recent work showed the production of a diversity of proteins in *A. platensis*, including single domain antibodies for oral delivery, there remains a need for a modular, characterized genetic toolkit^4^. Here, we establish and characterize a genetic toolkit and reproducible method for the transformation of *A. platensis* and engineer this bacterium to produce acetaminophen as proof-of-concept for small molecule production in an edible host from CO_2_, H_2_O, and light. This work opens *A. platensis* to the wider scientific community for future engineering as a functional food for nutritional enhancement, modification of organoleptic traits, and production of pharmaceuticals for oral delivery.

## Introduction

*Arthrospira platensis*, commonly referred to and sold commercially as spirulina, is an edible, non-heterocystous, filamentous cyanobacterium of the order *Oscillatoriales*, with species of this genus having been used as traditional food sources in Mexico and Chad for centuries^5,6^. *A. platensis* is considered safe for human consumption and is designated as Generally Recognized as Safe (GRAS) by the U.S. Food and Drug Administration (GFA GRN No. 417); it has exceptionally high protein content relative to plants and eukaryotic microalgae^1^, and is grown commercially as a food, pigment source, and animal feedstock^1,6^. Given its enticing food properties and established commercial growth, as well as applications driven by a photosynthetic metabolism, there is keen interest in genetically modifying this bacterium for enhanced nutrition, modified organoleptic traits, improved growth properties, and pharmaceutical production^1,2,4^.

While there are clear applications for sustainable biomanufacturing on Earth, the edible nature and photosynthetic metabolism of *A. platensis* makes it well-suited as an alternate food source and microbial host for on-demand pharmaceutical production for *in situ* resource utilization (ISRU)-based missions for human space exploration and habitation^7–9^. As humans delve deeper into space via the Moon and eventually to Mars and beyond, the need to reduce mission launch cost, reduce risk, and anticipate pharmaceutical degradation induced by radiation becomes paramount, further increasing the need for ISRU-driven food and pharmaceutical production^3,7,10–13^. NASA and ESA have both established the utility of this organism for space exploration via their biological life support projects, CELSS and MELISSA, respectively, which used *A. platensis* to photoautotrophically recycle NO_3_ ^-^ and CO_2_ for production of O_2_ and biomass for astronaut consumption^14,15^. Expanding upon its established life support properties to further produce functional foods for nutrition and pharmaceutical delivery will further increase the utility of this organism for deep space missions.

*A. platensis* has historically remained recalcitrant to the development of genetic tools and a reproducible transformation method^16–19^. Potential reasons range from its multicellularity, polyploidy, and motility, which combine to inhibit colony formation on plates, and a genome that encodes a high number of restriction modification systems of types I-IV in addition to several CRISPR/Cas systems^20,21^. However, in a landmark study, Jester *et al*. (2022) recently engineered *A. platensis* for antibody production targeting campylobacter infections, and showed that their orally-delivered strain prevents disease in mice and is safe for human consumption^4^. While this work provided a transformation method, the reported toolkit was used to express native operons or single, exogenous proteins using one of two endogenous, uncharacterized promoters, and is therefore limited in scope for the expression of multi-gene pathways to produce exogenous small molecules. Thus, a robust, characterized genetic toolkit that allows for the expression of exogenous multi-gene pathways in *A. platensis* remains unavailable. This limitation, combined with the general difficulty of performing genetic manipulations in this bacterium, have so far prevented *A. platensis* from being engineered for exogenous small molecule production.

Here, we develop two reproducible transformation methods based on electroporation and natural competence, use these methods to develop and characterize a genetic toolkit for downstream pathway engineering, and engineer *A. platensis* to produce acetaminophen as proof-of-concept for small molecule production derived from CO_2_, H_2_O, and light in an edible host for bioavailable, oral drug delivery.

## Results

### Genetic engineering of *Arthrospira platensis*

We initially developed an electroporation method inspired by the patent literature^22^ (see Methods for details). Due to its filamentous and motile nature, *A. platensis* does not form colonies on plates, and we were unable to obtain filament or cell counts. Thus, we were unable to quantify plating or transformation efficiency, and were therefore dependent on the binary output of growth or no growth, as has been observed in *Phormidium lacuna*^23^. Based on the predictions of Nies *et al*. (2020), we confirmed natural competence in *A. platensis*, and established this method as our standard transformation procedure.

Cyanobacteria have a wide range of genome copy numbers per cell^24^, with reported copy numbers of the *Oscillatoriales* ranging from 20-90 copies per cell in *Phormidium lacuna*^23^ to nearly 700 in *Trichodesmium erythreaum* IMS 101^25^. For all transformations resulting in biomass under selection, we observed that transformant biomass encoded the native locus in addition to the exogenous DNA (data not shown), suggesting an initial integration into the genome without full segregation through the genome and/or filament. Transformant biomass was subcultured until no native locus could be detected via PCR and/or genome sequencing. This took approximately 3-4 months on agar plates and 2-3 months in liquid culture (data not shown). Jester *et al*. (2022) reported a segregation time of 8-10 weeks for *A. platensis*^4^, further supporting our observations that exogenous DNA must be selected over long timeframes through a polyploid genome and multi-cell filament.

We established a series of working parts in *A. platensis* NIES-39: one antibiotic resistance marker, three genomic loci for insertion of exogenous DNA, four endogenous and five exogenous promoters, and one fluorescent reporter (Supplemental Table 1). All parts were integrated into a series of suicide vectors for homologous recombination with the genome (Supplemental Table 2). To screen multiple loci as genomic integration sites, we chose neutral site I (NSI; NIES39_Q01230), a commonly used site in other cyanobacteria for insertion of exogenous DNA^26^, along with two loci we expected to be involved in motility to potentially create non-motile strains to aid in plating and genetics.

Twitching motility in cyanobacteria is dependent on Type IV pili and is light regulated resulting in phototaxis towards or away from light sources^27,28^. Filamentous cyanobacteria from the *Oscillatoriales* and certain species of the *Nostocales* employ surface-dependent gliding motility, with the oscillin protein from *Phormidium uncinatum* being essential for this type of motility^27,29^. We targeted *pilA* (NIES39_C03030), a central component of the Type-IV pilin system, and an oscillin homolog (NIES39_A01430) as integration sites for exogenous DNA and motility ablation. We knocked out these three loci with the codon optimized version of *aadA, aadA*.*co*, which encodes for streptomycin/spectinomycin (Sm/Sp) resistance (Fig. 1; Supplemental Table 2). The ΔNSI::P*pilA*-*aadA*.*co*, Δoscillin::*aadA*.*co*, and Δ*pilA*::*aadA*.*co* strains were confirmed by whole genome sequencing (Fig. 1A). Of note, Δ*pilA*::*aadA*.*co* mutants display a “star” morphology when plated on agar, while Δoscillin::*aadA*.*co* mutants display a “string” morphology (Fig. 1). This implies that both genes are involved in motility and/or colony formation on plates.

**Figure 1.**
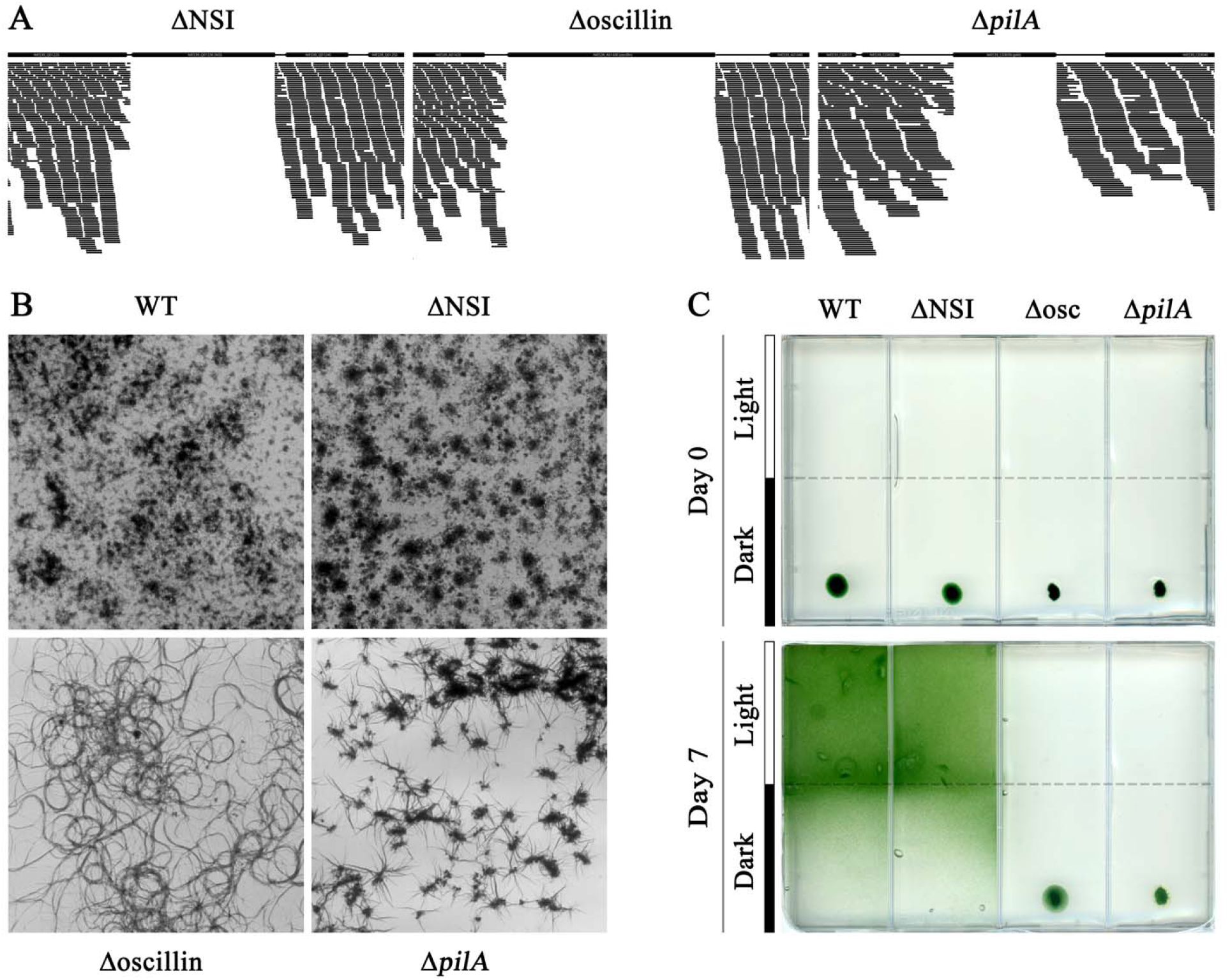
Confirmation and phenotyping of deletion mutants at three genomic loci. (**A**) Downsampled Illumina coverage for each genomic region that was swapped for *aadA*.*co*. (**B**) Images showing colony morphology on agar plates for the WT and each deletion strain. (**C**) Images depicting colony position on agar plates at Day 0 (top) and Day 7 (bottom) for the WT and each deletion strain. Strains were plated in the dark, and light was available to the upper half of the plate.

Deletion of the NSI locus does not result in morphological or growth differences between the wild type (WT) strain as expected (Fig. 1). Light-based motility assays showed that the WT and ΔNSI::P*pilA*-*aadA*.*co* strains moved into the illuminated area of the plate within seven days, while the Δoscillin::*aadA*.*co* and Δ*pilA*::*aadA*.*co* strains remained in the shade where they were initially spotted (Fig. 1C). While the Δ*pilA*::*aadA*.*co* strain did not expand beyond its initial spot, the Δoscillin::*aadA*.*co* mutant moved in a small halo around the initial spot, indicating that motility is severely limited by, but not entirely dependent on, the loss of oscillin. This indicates that both *pilA* and oscillin genes contribute to phototaxis in *A. platensis*.

### Characterization of promoter strengths using a fluorescent reporter system

To test an expression system in *A. platensis*, we designed a series of vectors (Supplemental Table 2) that resulted in mutant strains at the *pilA* locus that had one of three native, strong promoters (P_*psaA*_, P_*cpcB*_, or P_*rbcL*_)^30^ or one of five J23-series promoters driving the expression of codon optimized *eYFP* (*eYFP*.*co*) downstream of the native *pilA* promoter driving the expression of *aadA*.*co* followed by the native *aadA* terminator (Fig. 2A). The fluorescent output of each strain was measured, showing orders of magnitude higher fluorescence in the eight reporter strains than in the WT (Fig. 2B). The J23-series promoters displayed a 4-fold dynamic range in *A. platensis* compared to a 120-fold dynamic range in both *Synechococcus elongatus* UTEX 2973 and *Synechocystis* sp. 6803^31^. Although methodological differences between these studies may explain some of this difference, these results highlight the need for further development and characterization of *A. platensis* specific genetic parts. Fluorescence was manually checked under a fluorescence microscope to confirm expression of eYFP.co in the reporter strains; no fluorescence was detected in the WT strain (Fig. 2C).

**Figure 2.**
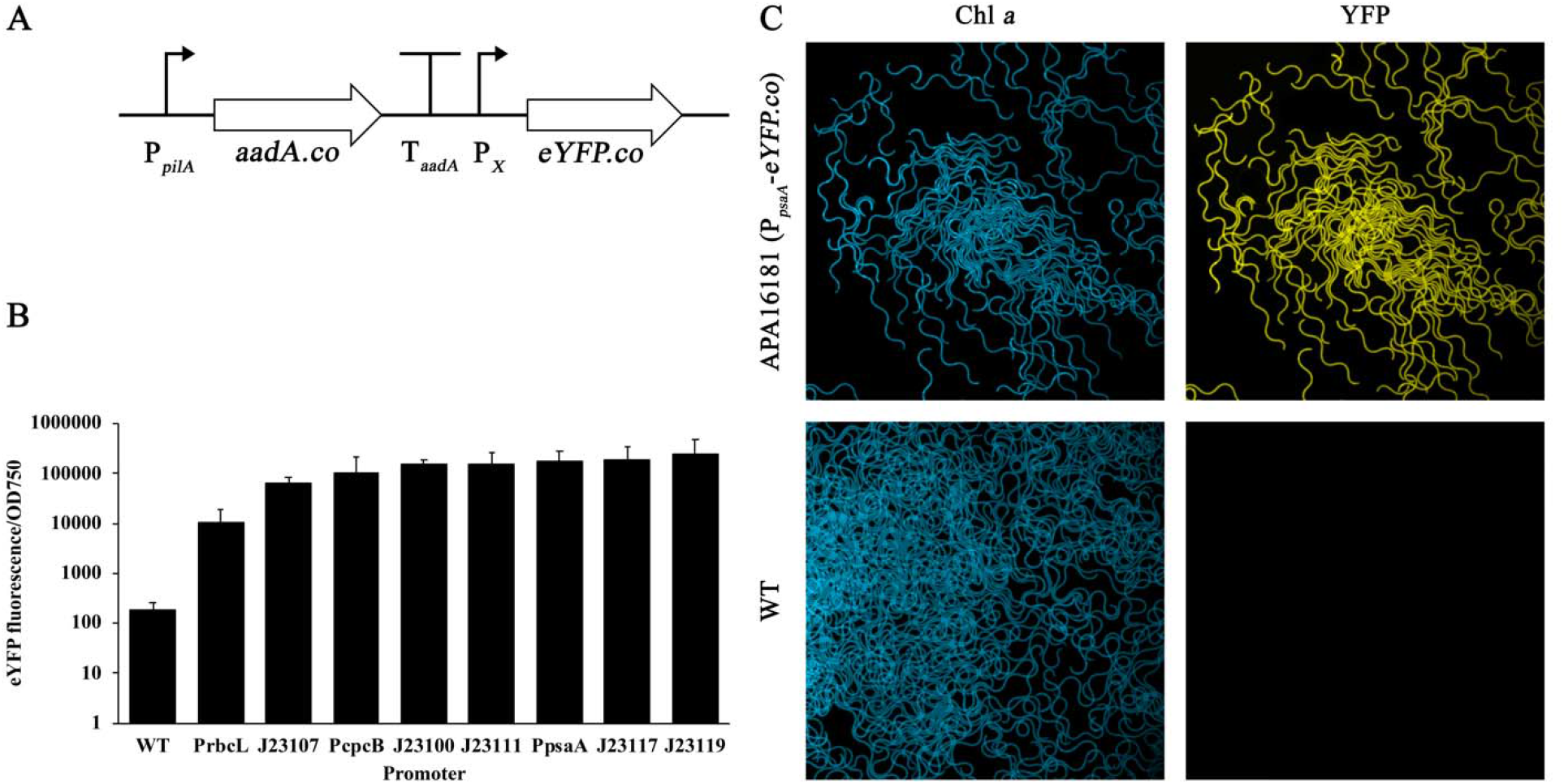
Fluorescent reporter and promoter characterization. (**A**) Schematic of the mutant genotypes for *eYFP*.*co* expression at the *pilA* locus: P_*pilA*_ drives expression of *aadA*.*co*, which is terminated by T_*aadA*_. Downstream, one of three native promoters drives expression of the *eYFP*.*co* gene. (**B**) Relative promoter strength for three native *A. platensis* promoters and the five J23-series promoters. Error bars represent standard deviations of technical replicates. (**C**) Fluorescence microscopy of the Δ*pilA*::*aadA*.*co*-P*psaA*-*eYFP*.*co* and WT strains showing Chl *a* autofluorescence of filaments under excitation at 587 nm, and YFP fluorescence under excitation at 485 nm.

### Acetaminophen production in *A. platensis*

For the biosynthesis of acetaminophen (APAP; Fig. 3A), chorismate is converted into para-aminobenzoic acid (PABA) by the multienzyme complex PabABC, composed of aminodeoxychorismate synthase (pabAB) and 4-amino-4-deoxychorismate lyase (pabC). PABA is converted into 4-aminophenol by the 4-aminobenzoate hydrolase (4ABH) from *Agaricus bisporus*, which is subsequently converted to APAP by N-hydroxyarylamine O-acetyltransferase (NhoA) from *Escherichia coli*^32^. A series of 38 suicide vectors encoding variations on the synthetic APAP biosynthesis pathway with codon optimized genes (Fig. 3A) were designed and screened in *A. platensis* (Supplemental Table 3). Of these, 23 resulted in transformant biomass that displayed the WT morphology under selection at 1 µg/ml Sm/Sp; no transformants for any vector could be recovered at higher selection strengths.

**Figure 3.**
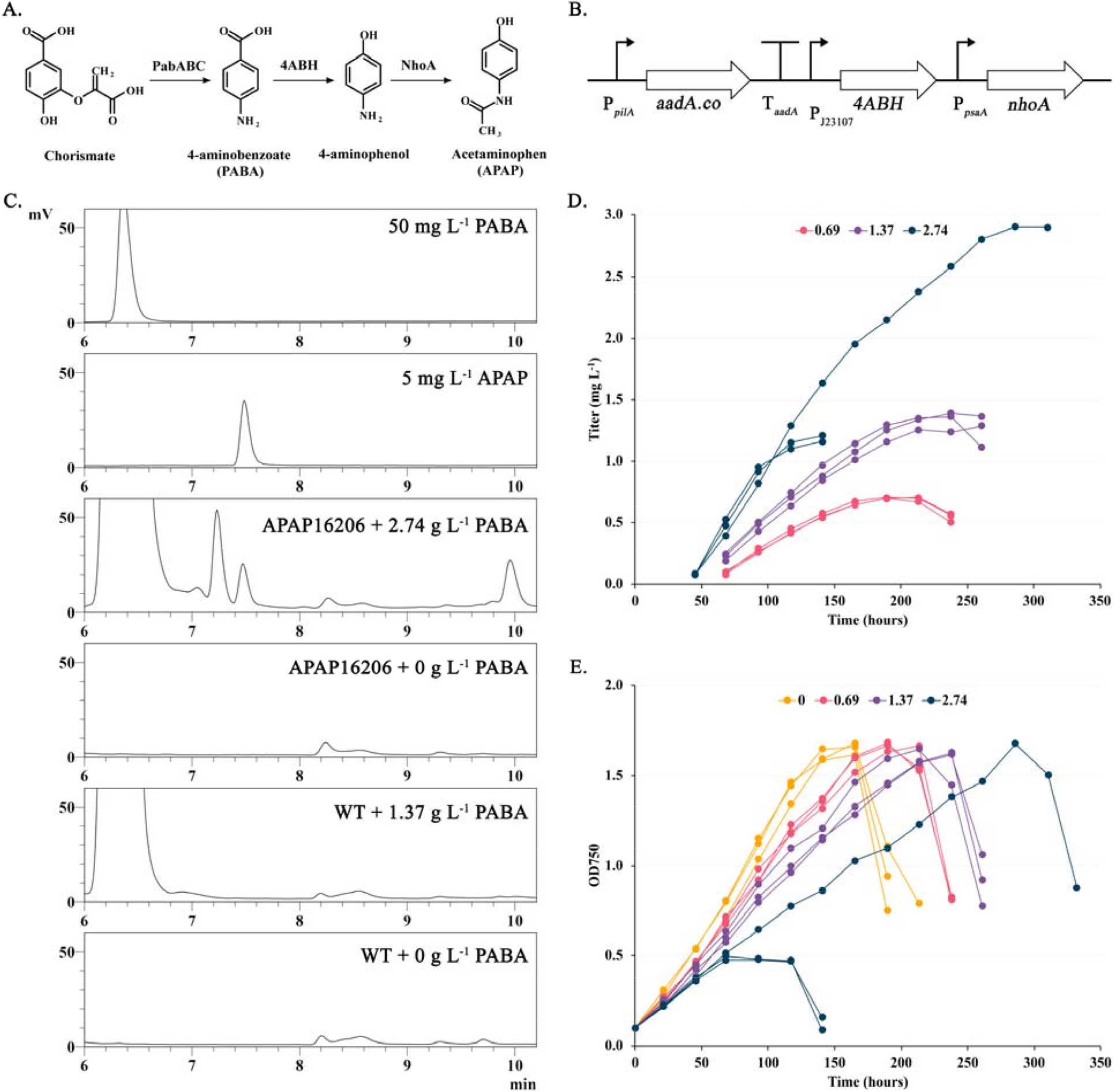
Figure 3. Acetaminophen (APAP) production in *A. platensis* APA16206. (A) The pathway for APAP production from para-aminobenzoic acid (PABA) uses the enzyme 4ABH to convert PABA to 4-aminophenol, which is then converted to APAP by NhoA. (B) Schematic for the APAP production pathway at the *pilA* locus in APA16206. (C) HPLC chromatograms showing peaks eluting between 6 and10.2 min for PABA and APAP standards along with APA16206 and WT strains with and without PABA supplementation. (D) APAP titer over time in APA16206 when supplemented with 0.69, 1.37, and 2.74 g L^-1^ PABA. (E) Growth of APA16206 with and without PABA supplementation at 0, 0.69, 1.37, and 2.74 g L^-1^.

For every transformation, a negative control was run to ensure that the selection killed WT cells, and pJMH024 was transformed as a positive control to ensure that cells were transformable. PCR screens of genomic DNA isolated from transformant biomass indicated the presence of *pilA, aadA*.*co, 4ABH*, and *nhoA* (data not shown), suggesting an initial integration of the suicide vector into the genome. However, no transformant biomass resulting from any of these vectors grew after subculturing into the same growth conditions. The only way we could recover a fully segregated strain, APA16206 (Fig. 3B), was under 16h light:8h dark cycles illuminated with ∼20-30 µmol m^-2^ s^-1^ (µEi) of photosynthetically active radiation (see Methods for details) for three months until star morphology appeared. Tufts of star morphology were subcultured into fresh selective medium under the same conditions. Upon reaching mid to late log phase, biomass was subcultured and grown under 24h light illuminated with ∼70-100 µEi. Biomass was regularly subcultured under these conditions upon reaching mid to late log phase until fully segregated (Supplemental Fig. 1).

As PABA supplementation was previously shown to increase APAP titer^32^, we screened APAP production in the WT and APA16206 strains with and without the addition of the precursor compound, PABA (Fig. 3C-E; Supplemental Fig. 2). The WT strain did not produce APAP under any condition, while the mutant produced APAP only when PABA was added to the medium (Fig. 3C). The APAP titer and production rate in APA16206 were dependent on the amount of PABA supplied, with titer and rate increasing with increasing PABA concentrations. The maximal titer and rate were 2.9 mg L^-1^ and 0.018 mg L^-1^ hr^-1^, respectively, when supplied with 2.74 g L^-1^ PABA. Growth rate was slowed by increasing PABA concentration, but did not impact the final biomass yield, except for two of the replicates supplemented with 2.74 g L^-1^ PABA, which prematurely reached stationary phase and died (Fig. 3E). The APAP production rate appeared dependent on the growth rate.

## Discussion

Here we independently establish the reproducible transformation of *A. platensis* via electroporation and natural competence, characterize an initial genetic toolkit for pathway engineering, and produce APAP as a proof-of-concept pharmaceutical for oral delivery. Our methods help unlock this bacterium for future engineering efforts for sustainable biomanufacturing on Earth and ISRU-based food and pharmaceutical production in space by providing a reproducible blueprint. Building on recent work^4,23^, we help establish genetic tractability in the *Oscillatoriales*, an order of cyanobacteria that have been historically understudied through genetic methods. Members of this order, such as the diazotrophic genus *Trichodesmium*, are major contributors to the biological nitrogen cycle.

While our open-source genetic toolkit and proof-of-concept biosynthesis of APAP is an important step for metabolically engineering *A. platensis*, continued improvement in the genetic tools available is paramount to decrease the design-build-test-learn cycle time in this bacterium. Crucial to this endeavor is the development of new promoters to overcome the limited dynamic range we observed, which may be achieved through the characterization of alternate native promoters, synthetic promoters engineered from native promoters, or the establishment of inducible promoters^33^. Operationalizing a CRISPR/Cas system^34^ would provide a locus-independent counterselection that may decrease the segregation time, as the Cas enzyme would be expected to cut all, or most, genome copies present in the cell,, and would allow for installation of more complex pathways. Developing self-replicating vectors for the expression of pathways or a CRISPR/Cas system would bypass the need for genome integration, and allow for rapid screening of pathway topologies and simpler CRISPR/Cas-based genome engineering.

APAP is the first reported exogenous small molecule produced in *A. platensis* and provides a milestone in the long-term efforts to genetically engineer this organism. Given that a dose of APAP is 325 mg, our maximum titer of 2.9 mg/L would require ∼112 L for a single dose, assuming complete bioavailability. Thus, higher titer will need to be achieved to make manufacturing a dose of APAP feasible if oral delivery via spirulina is the goal. To this end, there are several routes that should be explored.

Of the 38 APAP pathway topologies we transformed into *A. platensis*, we could only recover one fully segregated mutant that, in turn, could only be recovered under day/night cycles. This was in stark contrast to the creation of other mutants reported here, which proceeded without difficulty, and suggests that enzyme toxicity from 4ABH and/or NhoA, due to their products or the enzymes themselves, inhibited integration and/or segregation of the pathway into and through the genome. Several additional, unknown chromatography peaks were observed in APA16206 when PABA was supplied (Fig. 3C), and may contribute to the observed pathway toxicity. These unknown peaks eluted at similar times as PABA and APAP, implying a similar chemical structure. These peaks may be due to enzyme promiscuity of 4ABH and/or NhoA, or from APAP degradation products; these peaks are not from the intermediate 4-aminophenol as the standard for this molecule produced two peaks at elution times of 2.9 and 3.3 min (data not shown). If these compounds are produced from enzyme promiscuity, engineering a strain to produce only APAP would likely increase the titer of APAP by avoiding flux of PABA into these accessory compounds. This could be done through enzyme engineering, removing biosynthesis pathways from the genome for substrates that these enzymes may promiscuously convert, or screening 4ABH and/or NhoA homologs for improved selectivity towards APAP production. Decreasing the potential pathway toxicity through use of weaker promoters, enzyme scaffolding, or enzyme engineering for higher product selectivity may make this pathway more engineerable for future iterations towards a higher titer.

PABA was not detected in either strain without PABA supplementation. This may be due to natively low flux through PABA or a high rate of PABA turnover. However, while we were able to identify a homolog for PabAB (NIES39_E04160), we were unable to identify a homolog for characterized PabC proteins^35–37^ in the *A. platensis* genome. While there may be an uncharacterized PabC, there is a chance that PABA is not natively produced by *A. platensis*, which would be unexpected as PABA is a precursor for folate. Establishing and/or increasing endogenous PABA biosynthesis could increase APAP titer, or, crucially, allow for production without PABA supplementation. As the APAP yield from supplied PABA was ∼0.1%, a sufficient intracellular production rate of PABA may be easily achievable to reach APAP production rates observed with supplementation. This may also alleviate the observed toxicity from PABA when supplied at high extracellular concentrations (Fig. 3D). Engineering *A. platensis* for sufficient PABA production would also remove the economic burden of PABA supplementation for scale-up.

The genetic toolkit and methods we provide here pave the way for the diversification of commodity chemicals and biologics to be produced in *A. platensis* beyond APAP and the expressed proteins of Jester *et al*. (2020)^4^. Decisions on which molecules to produce would benefit from a combination of technoeconomic and life cycle analyses targeting high value products such as pharmaceuticals and cannabinoids. This work further opens *A. platensis* as an engineerable, functional food. Modification of organoleptic traits and nutritional properties may lead to more palatable and nutritious strains for improved consumption by humans or for animal feedstocks. Improved genetic tools along with host optimization; for example, through the removal of restriction modification systems, genome copy number control, and morphological control, may further improve the engineerability of this organism. The present work is thus an important step towards realizing the potential of an industrially grown, edible, photosynthetic organism for the oral delivery of pharmaceuticals and nutritional compounds.

## Online Methods

### Growth and transformation of *Arthrospira platensis*

A list of strains used in this study can be found in Supplemental Table 2. WT *A. platensis* NIES-39, obtained from the NIES culture collection, was grown in Zarrouk’s Medium (ZM)^38^ at 30°C under ∼70-100 µmol photons m^-2^ s^-1^ (µEi) of photosynthetically active radiation produced by cool, white LEDs (compact LED bars from Photon Systems Instruments controlled by their LC 200 light controller) with shaking at 200 rpm unless stated otherwise. Working streptomycin/spectinomycin (Sm/Sp) concentrations ranged from 1-100 µg/ml.

For electroporations, WT *A. platensis* NIES-39 cells were grown to early log phase (OD_750_ ∼0.2-0.3), washed three times with room temperature, sterile water and concentrated to 150 µg Chl *a* ml^-1^ in sterile water. Chl *a* concentration was measured based on the methods of Meeks and Castenholz (1971)^39^. For each electroporation reaction, 400 µl concentrated cells were mixed with 10 µg plasmid on ice, transferred to a cold 2 mm electroporation cuvette, and electroporated at 600 V with a time constant of ∼6-7 ms^-1^ on a BTX Gemini electroporator. Cells were recovered in a final volume of 1 ml ZM, and grown overnight (∼15-20 hours) at 30°C under ∼20-30 µEi with shaking. The following day, cells were washed once with ZM + Sm/Sp^1^, and plated onto selective 1.5% Bacto agar (BD #214010) plates or into selective liquid ZM in multiwell (96-, 24-, 12-, and/or 6-well) plates. Biomass was selectively passaged onto plates and into liquid media, and monitored via PCR and/or Illumina sequencing until fully segregated.

For natural transformations, WT *A. platensis* NIES-39 cells were grown to early log phase (OD_750_ ∼0.2-0.3) and concentrated to 150 µg Chl *a* ml^-1^ in ZM. 100 µl concentrated cells were mixed with 2.5 µg plasmid DNA, and incubated for 45 min at room temperature. Cells were recovered in a final volume of 1 ml ZM, and grown overnight (∼15-20 hours) at 30°C under ∼20-30 µEi with shaking. Recovered cells were added to 99 ml ZM + Sm/Sp (1-10 µg/ml) in 250-ml flasks, and incubated under ∼20-30 µEi until growth appeared. Biomass was subcultured under increasing concentrations of Sm/Sp (up to 100 µg/ml) until fully segregated as monitored by PCR.

### Genomic DNA Extraction

gDNA was extracted as follows: 1-ml aliquots of culture were harvested by centrifugation. Pelleted cells were resuspended in 300 µl ELB (20 mM Tris-HCl, 2 mM Na-EDTA, 1.2% Triton-X). Cell suspension was incubated at room temperature for 30 min with 2,500 units of Ready-Lyse lysozyme (Lucigen #R1804M) and 200 µg RNase A (Qiagen #19101). An additional 2,500 units of lysozyme were added and incubated for 30 min at room temperature. 15 µl of Proteinase K (20 mg/ml; Qiagen #19157) were added to the suspension and incubated at 65°C for approximately one hour. 400 µl phenol:chloroform:isoamyl alcohol (25:24:1 v/v; Sigma #P3803) were mixed with the cell suspension, and centrifuged for 5 min at 21,130 *x* g. 30 µl of 3M sodium acetate + 0.1M EDTA were added to the aqueous phase, followed by 660 µL cold 100% ethanol, and centrifuged for 15 min at 21,130 *x* g. Supernatant was removed, and the pellet was washed once with 1 mL 70% EtOH. Following removal of the wash, the pellet was air dried for 5 minutes prior to resuspension in nuclease-free TE buffer (Thermo Scientific #J75793).

### Next Generation Sequencing

Library preparation and 250 bp paired-end sequencing on an Illumina HiSeq 4000 was completed by the QB3 Genomics, UC Berkeley, Berkeley, CA, RRID:SCR_022170.

### Data analysis

Raw Illumina NovaSeq sequencing reads were processed as follows. Burrows-Wheeler Alignment tool (BWA)^40^ was used to index the reference genome AP011615, align sequencing reads and generate a sam file. Samtools^41^ was used to convert the data to a bam file and then sort and index the alignments. Picard’s DownsampleSam (https://broadinstitute.github.io/picard/) was used to downsample the BAM files to ∼10% of the total depth for visualization in Geneious. Geneious was used to visualize the alignments and generate figures.

### Motility assays

ZM solidified with 0.5% agar was poured into each lane of a 4-well plate. Cultures of each strain to be assayed were inoculated to a starting OD_750_ of 0.05 and allowed to grow until mid-log phase (OD_750_ = 0.5 – 0.7). A volume of each culture equal to 750 μL OD_750_ = 1.0 was pelleted by centrifugation and the supernatant was removed. The cell pellets were collected and dispensed at one end of an agar lane in the 4-well plate. The plate was sealed and imaged immediately. The plate was moved to a 30°C room and placed directly under cool, white LEDs at a flux of 50 µEi. The half of the plate containing the cell pellets was placed in a protective covering to prevent direct exposure to light and encourage growth to the far end of the plate. The plate was imaged every 24 hours for 7 days.

### Vector design and synthesis

A list of vectors can be found in Supplemental Table 2. All genes were codon optimized, purged of the restriction modification (RM) sites (see below), synthesized, and cloned by Genscript Corp. The remaining parts were synthesized and cloned by Genscript Corp. The purged RM sites can be found in Supplemental Table 4. This list was compiled from literature and homology searches^20,42,43^.

### Fluorescence microscopy

Mid-log phase cultures were imaged on a Zeiss Observer D1 fluorescence microscope. Chlorophyll autofluorescence was imaged under excitation at 587 nm and emission at 610 nm and eYFP fluorescence was measured under excitation at 495 nm and emission at 527 nm. Images were falsely colored using Zeiss Zen Pro version 3.5.

### Quantification of promoter strengths

*A. platensis* WT and mutant strains were each grown in eight wells of triplicate 96-well plates under ∼20-30 µEi without shaking at 30°C. OD_750_ and eYFP fluorescence (excitation at 485 nm and emission at 535 nm) were measured daily on a Tecan Spark M10 plate reader, and these values were normalized to blank measurements.

### HPLC

Acetaminophen (APAP) and para-aminobenzoic acid (PABA) were quantified on a Shimadzu Prominence HPLC system equipped with a photodiode array detector for UV detection. Compounds were separated on a Zorbax StableBond Plus C18 5 µm with detection at 240 nm. The elution protocol was based on a gradient method using (A) 0.1% phosphoric acid and (B) methanol as follows: 0 min - 95% A, 5% B; 15 min 100% B; 16-20 min 95% A, 5% B at a flow rate of 1 ml min^-1^.

For sample preparation, 1 ml culture was transferred to a BeadBug bead beating tube (Millipore Sigma #Z763748) and vortexed on high for 10 minutes. Samples were then centrifuged for 10 min at 14k rpm. 800 µl supernatant were filtered through 13 mm 0.2 µm PVDF syringe filters (Pall #4406). Standards were prepared in volumetric flasks using APAP (Sigma #A5000) and PABA (Sigma #A9878), and serially diluted in ZM. The 4-aminophenol (Sigma #A71328) standard was prepared similarly using methanol as the solvent.

## Acknowledgements

This work was supported by the Center for the Utilization of Biological Engineering in Space (CUBES, https://cubes.space), a NASA Space Technology Research Institute (grant number NNX17AJ31G) and by NIH S10 OD018174 Instrumentation Grant. We thank Kyle Sander, Aaron Berliner, Kelly Wetmore, Kelsey Hern, and Yolanda Huang for useful discussions and support.

## Author Contributions

J.M.H., J.M.S., and A.P.A. conceived the project. J.M.H., S.F., J.M.S., and A.P.A. designed experimental work. J.M.H., S.F., D.S., S.Y., and J.M.S. performed experimental work. J.M.H., S.F., J.M.S., and A.P.A. analyzed data. Y.C. developed the HPLC method. J.M.H., S.F., J.M.S., D.S.C., and A.P.A wrote the manuscript.

## Supplementary Data

Plasmid maps for plasmids listed in Supplementary Tables 2 and 3 are available at https://doi.org/10.6084/m9.figshare.20085284.

