## Supplemental Figures for "Acetaminophen production in the edible, filamentous cyanobacterium *Arthrospira platensis*"

5-Molecular Biophysics and Integrated Bioimaging Division, Lawrence Berkeley National

Laboratory, 1 Cyclotron Road, Berkeley, CA 94720, USA

6-Environmental Genomics and Systems Biology Division, Lawrence Berkeley National Laboratory, 1 Cyclotron Road, Berkeley, CA 94720, USA

**Supplementary Figures**

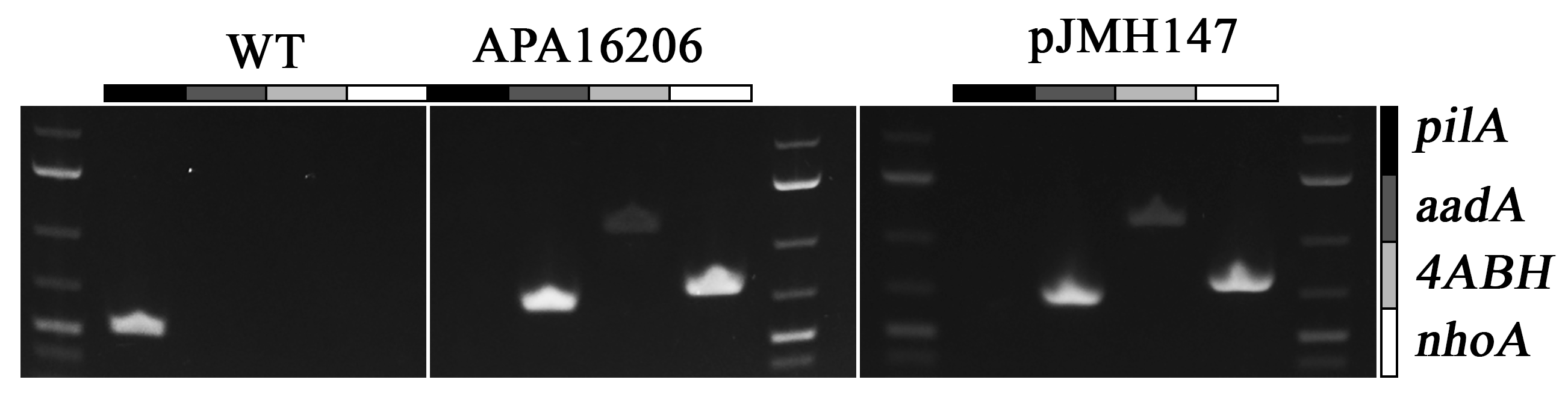

**Supplemental Figure 1.** Confirmation of mutant genotype in APA16206. PCR gels for amplification of *pilA*, *aadA*, *4ABH*, and *nhoA* in the WT and APA16206 strains along with the suicide vector used to create APA16206, pJMH147.

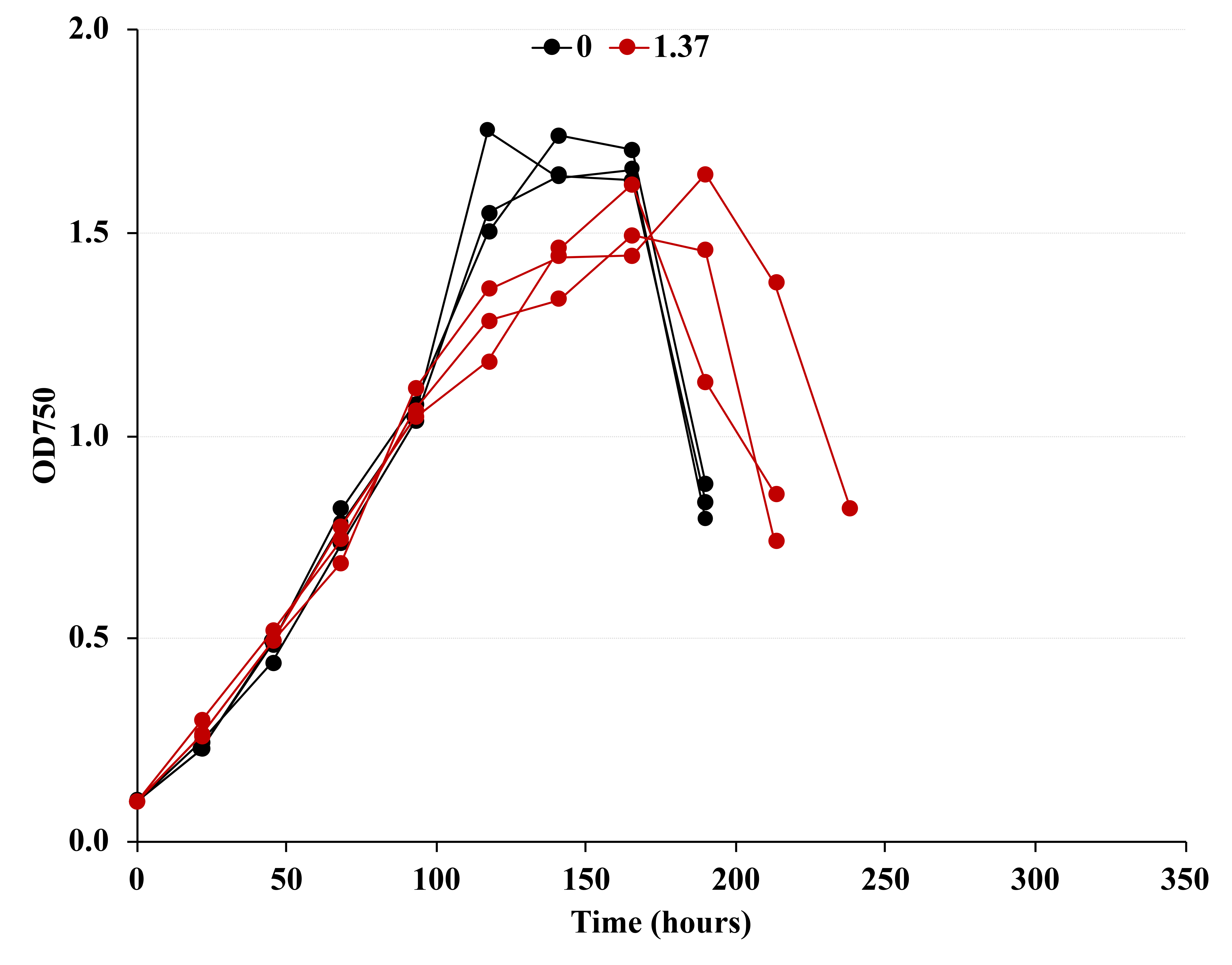

**Supplemental Figure 2.** Growth of the WT strain with (red) and without (black) PABA supplementation at 1.37 g L^-1^.
